# Data-driven discovery of canonical large-scale brain dynamics

**DOI:** 10.1101/2022.07.27.501789

**Authors:** Juan Piccinini, Gustavo Deco, Morten Kringelbach, Helmut Laufs, Yonatan Sanz Perl, Enzo Tagliazucchi

**Author notes:** Corresponding authors: Juan Piccinini, Enzo Tagliazucchi.

## Abstract

Human behavior and cognitive function correlate with complex patterns of spatio-temporal brain dynamics, which can be simulated using computational models with different degrees of biophysical realism. We used a data-driven optimization algorithm to determine and classify the types of local dynamics that enable the reproduction of different observables derived from functional magnetic resonance recordings. The phase space analysis of the resulting equations revealed a predominance of stable spiral attractors, which optimized the similarity with the empirical data in terms of the synchronization, metastability, and functional connectivity dynamics. For stable limit cycles, departures from harmonic oscillations improved the fit in terms of functional connectivity dynamics. Eigenvalue analyses showed that the proximity to Hopf bifurcations improved the accuracy of the simulation for wakefulness, while deep sleep was associated with increased stability. Our results provide testable predictions that constrain the landscape of suitable biophysical models, while supporting noise-driven dynamics close to a bifurcation as a canonical mechanism underlying the complex fluctuations that characterize endogenous brain activity.

## Introduction

Brain dynamics are often described as complex, displaying properties that are interposed between order and disorder^1-3^. These complex dynamics arise from two main factors: the properties of local population activity within each brain region and the mutual influences that these populations exert on each other^4,5^. Over the last few years, different kinds of models have been introduced to disentangle the contributions to whole-brain dynamics and their relationship with cognition and behavior^6-9^. By combining empirical data with simulated local dynamics, models of whole-brain activity have been applied to describe multiple physiological and pathological states, allowing to explore the potential mechanisms underlying different neurobiological phenomena, and offering the possibility of in silico assessment of external perturbations^8,10-12^. Generative models based on dynamical systems can also be used for data augmentation, in combination with deep learning or other methods from machine learning^13-15^. Importantly, whole-brain models are capable of furnishing concrete falsifiable hypotheses by virtue of their grounding in individualized empirical data^16^.

What properties should a computational model possess to accurately represent large-scale brain activity dynamics? A sufficient degree of biophysical detail is necessary to link the outcomes of the model with neurobiological variables of interest, such as axonal conduction delays, stimulation of neurotransmitter receptors, or changes in synaptic gating, among others^17-21^. However, biophysical realism does not guarantee that simulated brain activity will display the statistical properties measured in empirical data. For this purpose, it is important that models exhibit certain stereotyped behaviors capable of generating dynamics of sufficient complexity^1,2^. In other words, the equations of the model should display certain dynamical behaviors that can be better understood in terms of the topology of the phase space (i.e., the space of possible solutions) than in terms of the biophysical details of the model^22^. One example is noise-driven multistability, where stochastic fluctuations displace the state across a bifurcation, switching between two or more qualitatively different solutions, such as stable vs. dampened oscillations^23^. Thus, the core capacity of a model to capture whole-brain dynamics can be characterized by its repertoire of bifurcations and their classification. For instance, noise-driven models close to a bifurcation (such as the Stuart-Landau oscillator) have been extensively explored and characterized in recent publications^13,24-27^. Even though more realistic models offer advantages in terms of interpretability, they cannot escape the fact that most of the time, if not always, the model parameters must be posed next to a bifurcation to adequately reproduce empirical observables^22^.

The process of building and validating a whole-brain activity model usually begins with the hypothesis-driven proposal for the equations governing the local dynamics, followed by the exploration of parameter space to maximize the goodness of fit to the empirical neuroimaging data^6^. However, focusing on a particular set of equations could be too constraining, since the appropriateness of a model should be judged at a different level, namely by its capacity to reproduce certain stereotyped dynamics present in the empirical data^22^.

Here, we tackled this problem by following the inverse procedure: we first proposed very general equations, and then we fitted these local equations of motion to observables derived from functional magnetic resonance imaging (fMRI) data^28^, characterizing the resulting equations in terms of their attractors and their proximity to bifurcations. This procedure is data-driven and independent of specific model details, and its outcome can be interpreted as the set of canonical dynamics that are desirable in whole-brain activity models of fMRI recordings.

## Results

An overview of the procedure is presented in Fig. 1, with further details provided in the methods section. Briefly, we proposed local dynamics given by two equations, corresponding to variables x(t) and y(t) which were combined to form all possible polynomial terms with degree less or equal than C. Only variable x(t) represented the simulated brain activity signal; the other was a hidden variable necessary to endow the system with non-trivial dynamics. These equations were coupled by the connectome scaled by parameter G and included additive noise scaled by factor κ (following previous research, G was sufficiently small to allow a weak coupling approximation)^25^. Polynomial equations were chosen based on their generality, since it is known that other functions can be replaced by their low order polynomial approximation when investigating the normal form of different bifurcations^28,29^.

**Figure 1.**
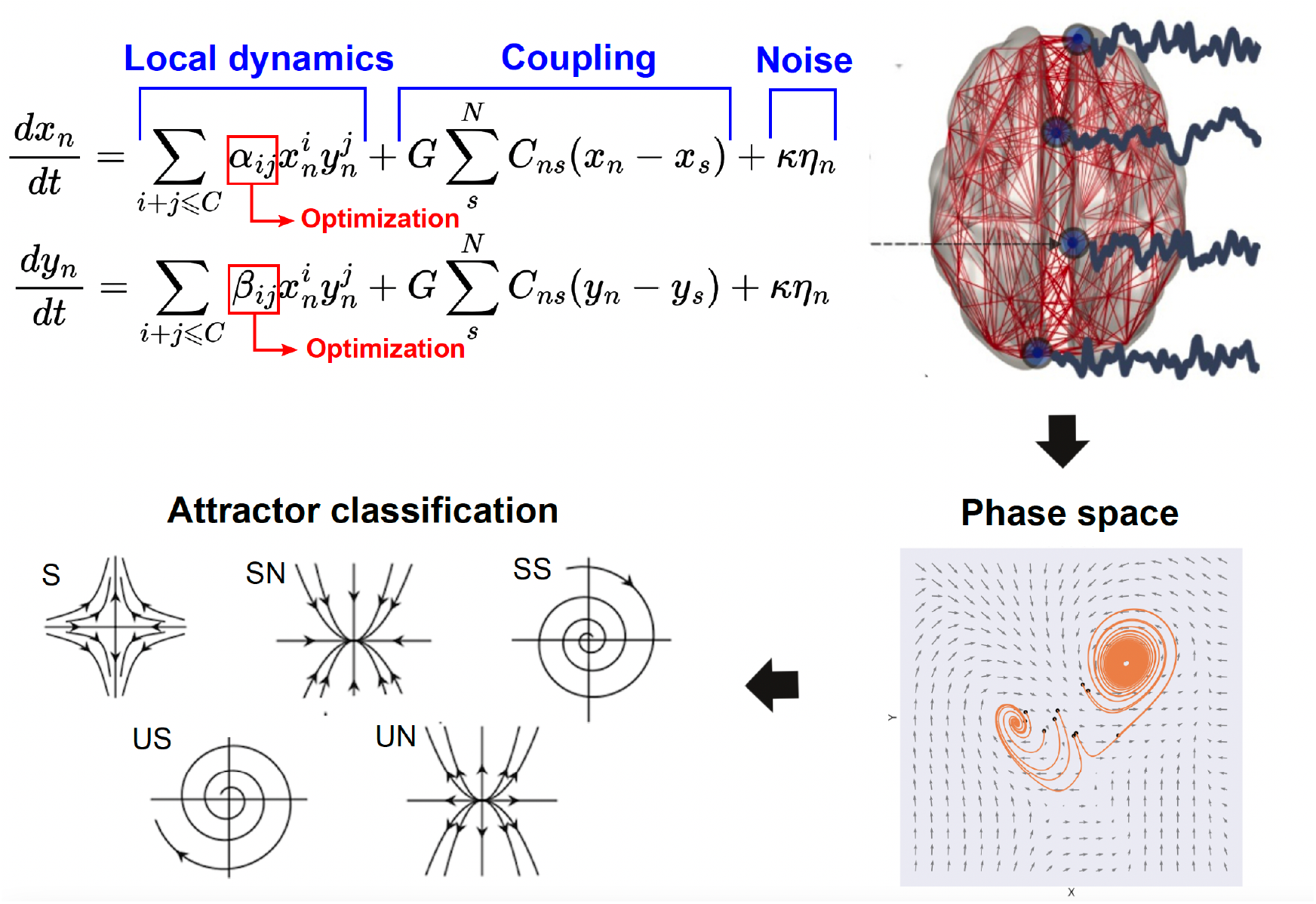
Procedure followed for the data-driven discovery of canonical whole-brain dynamics. Each iteration of the model consisted in local dynamics given by the two variables x, y combined in polynomial terms up to degree C with coefficients α_ij_, coupling by the connectome scaled by G, and noise scaled by κ. After the initial selection of G, the parameters α_ij_ were optimized to reproduce fMRI functional connectivity between all pairs of nodes. The optimal local dynamics can be characterized in terms of the 2D phase space of variables x, y, where different attractors can be identified and used to characterize the resulting dynamics.

We performed 1000 independent iterations of the optimization procedure, setting the degree of the polynomial C=5, which resulted in 42 free parameters to be determined by the stochastic optimization algorithm (genetic algorithm)^27^. Each iteration attempted to maximize a metric of similarity computed both for the simulated and empirical data (structural similarity index [SSIM] between the corresponding functional connectivity matrices)^30^. We opted to use the SSIM since it balances sensitivity to absolute (e.g. euclidean distance) and relative (e.g. correlation distance) differences between the functional connectivity matrices. After optimization, the resulting local dynamics were visualized in phase space, and numerical methods (i.e., analysis of the Jacobian matrix) were used to infer the presence of different attractors and the proximity to bifurcations (see the methods section for an overview of the classification criteria).

Considering the introduction of noise in the dynamics, we did not expect the optimization algorithm to converge to the exact same set of coefficients α_ij_ across all iterations. Instead, we focused on the statistical characterization of the optimal dynamics and their properties. Figure 2A presents the number of solutions with one, three and five fixed points in the phase space, with parameters optimized to match the functional connectivity matrix of awake individuals. A fixed point corresponds to a pair x, y where the derivatives 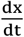 and 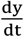 are both zero, so that dynamics starting at that point cannot change over time. We found that the most likely outcome consisted of a single fixed point, followed by three fixed points, with a comparatively small number of optimal equations presenting five fixed points. Next, we asked whether the number of fixed points impacted on the similarity to the empirical data, assessed by four independent fMRI observables and their associated goodness of fit metrics: 1-SSIM between empirical and simulated functional connectivity matrices, synchronization, metastability^31^ and the Kolmogorov-Smirnov distance between the empirical and simulated distributions of functional connectivity dynamics (FCD) values^25^ (see the methods section for a definition of these observables). The results shown in Figure 2B indicate that these metrics do not depend on the number of fixed points in the local dynamics.

**Figure 2.**
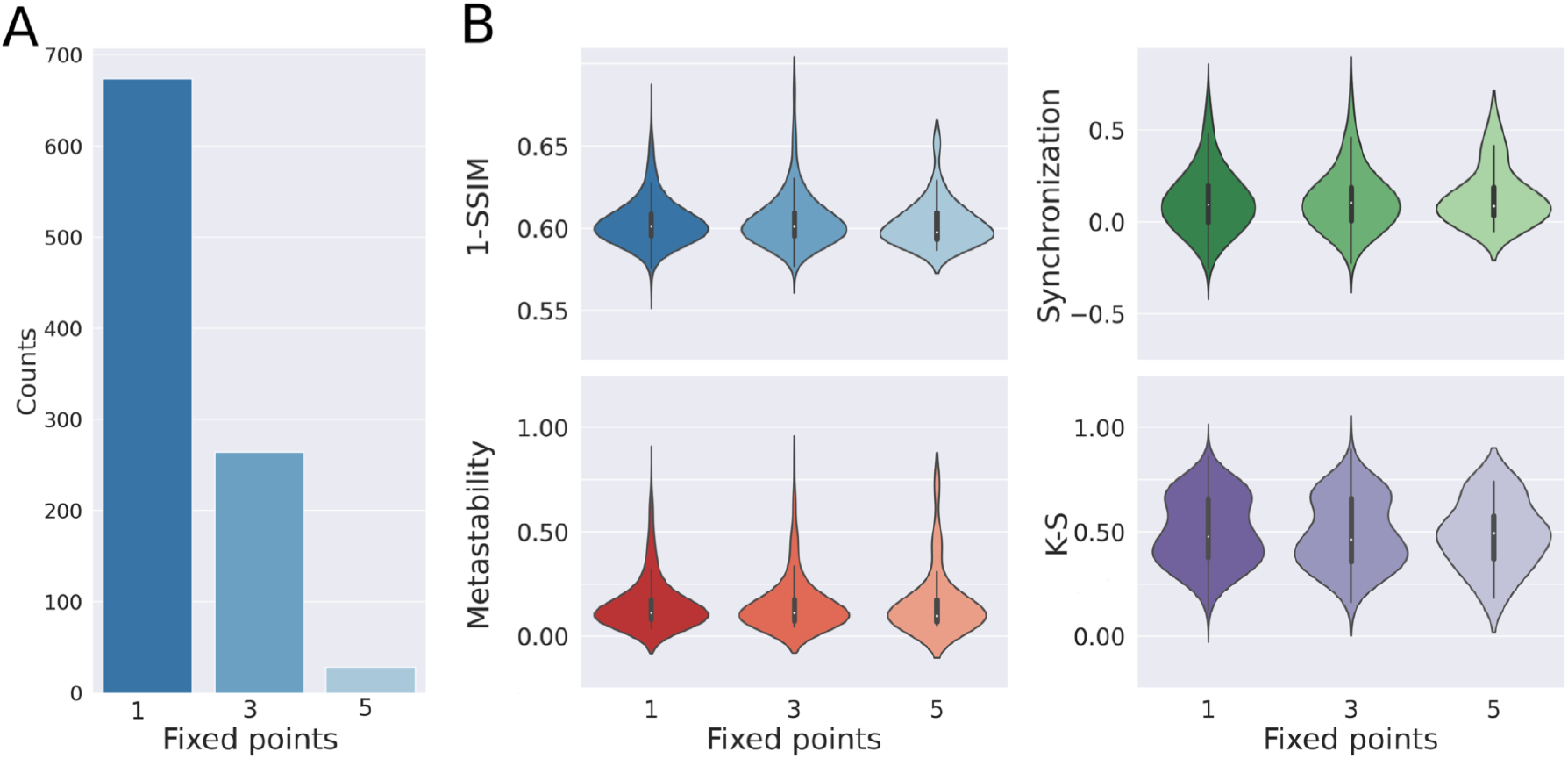
Local dynamics tend to exhibit a single fixed point, and the similarity between simulated and empirical dynamics is independent of the number of fixed points. A) Number of iterations resulting in one, three and five fixed points. B) Four different metrics computed after separating the solutions by the number of fixed points in the phase space. No differences were encountered when comparing local dynamics with different numbers of fixed points.

Next, we classified the isolated fixed points based on the analysis of the Jacobian matrix, among the following possibilities (see Fig. 1, “attractor classification”): stable node (SN), unstable node (UN), saddle node (S), stable spiral (SS) and unstable spiral (US). We found that all isolated fixed points were spirals, with a predominance of stable spirals (i.e., damped oscillations) (Fig. 3A, left). In the case of unstable spirals, all instances were surrounded by limit cycles, asymptotically leading to bounded oscillatory solutions. Only 2% of the optimal equations resulted in stable spirals surrounded by limit cycles, which have potential to display bistable oscillatory dynamics. Even though stable spirals appeared more frequently in the local dynamics, the goodness of fit metric 1-SSIM was comparable for both types of spirals (Fig. 3A, right). Both types of fixed points are exemplified in the phase portraits shown in Fig. 3B. Finally, panel C of Fig. 3 contains a scatter plot of the imaginary vs. real eigenvalues for each iteration, illustrating the separation between stable and unstable solutions given by the vertical line of null real eigenvalues.

**Figure 3.**
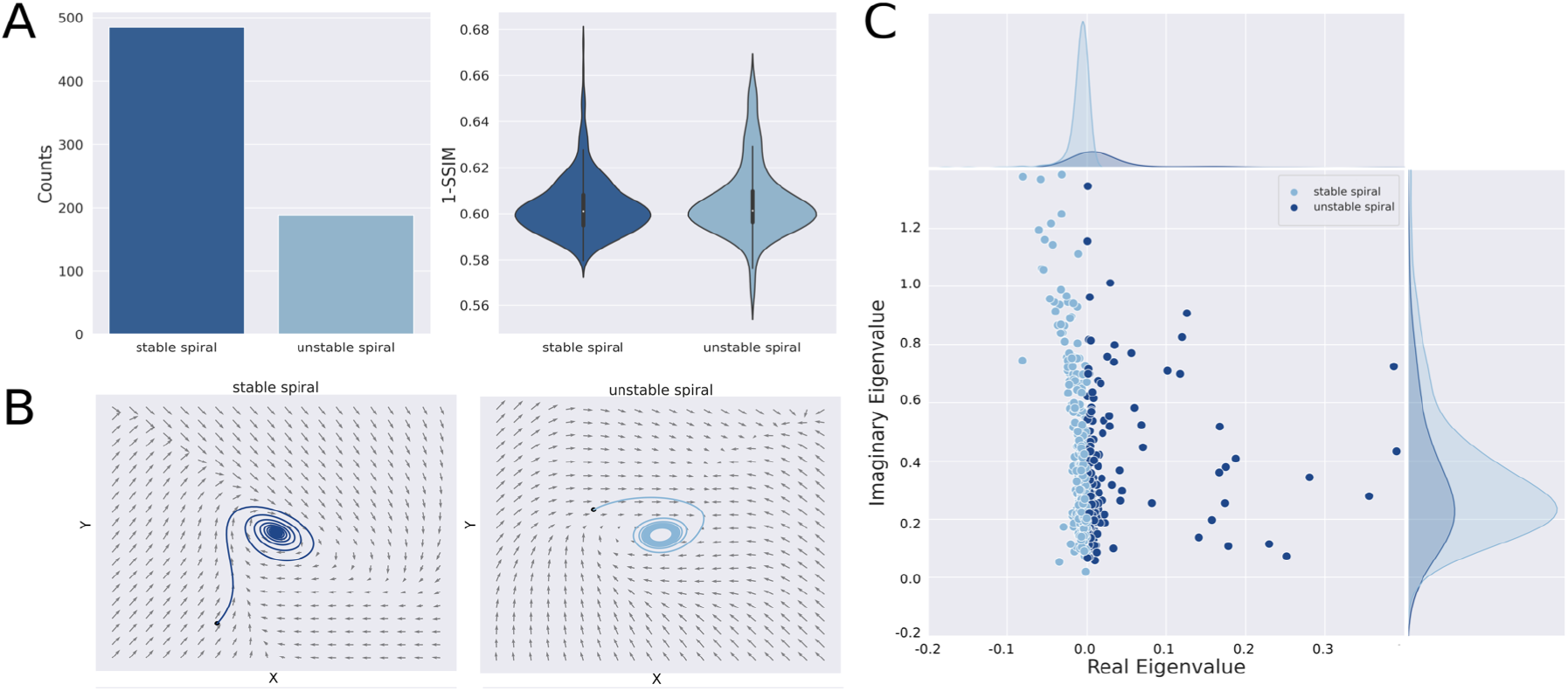
Stable spirals are most prevalent for local dynamics with one isolated fixed point. A) Left panel: Number of iterations resulting in local dynamics with stable and unstable spirals. Right panel: 1-SSIM for both types of local dynamics. B) Examples of phase spaces with each type of local dynamics. Note that the unstable spiral is surrounded by a limit cycle (attractor consisting of a periodic trajectory). C) Scatter plot of the imaginary vs. real eigenvalues of the fixed point, where each point corresponds to an independent iteration of the model. The vertical line of null real eigenvalues determines the stability of the spiraling solution.

Local dynamics with three fixed points were the second most probable outcome (Fig. 2A). We coded each possible combination using the above introduced abbreviations; for example, S-SN-SS identified local dynamics with a saddle node, a stable node and a stable spiral. The statistics for the case of three fixed points are shown in Figure 4. Here, the entries of the matrix indicate the number of times each possible type of fixed point labeled in the rows appeared in the optimal local dynamics specify in the columns. For example, the value 51 in the third row and third column indicates a total of 51 stable spirals within the combination S-SS-US, while the sum of all column values indicates the number of times the combination S-SS-US was found throughout the 1000 iterations. We note that several solutions were possible, yet these tended to be dominated by stable spirals and saddle nodes, with S-SN-SS being the most frequent combination, followed by S-SS-SS and S-SS-US. This suggests that dynamics still find their way to stable spiral attractors after being repelled by saddle nodes or unstable spirals. Overall, local dynamics where stable spirals appeared as part of the phase space were much more likely to be found than those containing other fixed points, in agreement with the findings obtained for isolated fixed points. Examples of trajectories for different combinations of attractors are shown in Fig. 4B.

**Figure 4.**
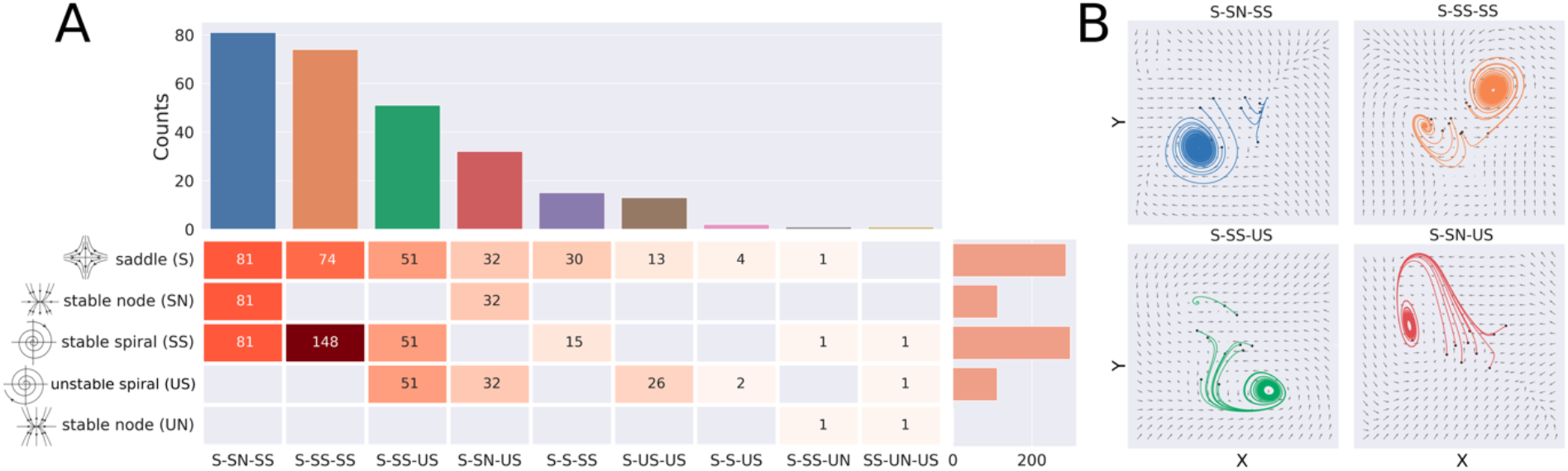
Saddle nodes and stable spirals are the most predominant for local dynamics with three fixed points. A) Matrix entries indicate the total number of fixed points (rows) that are present in a specific combination (columns). The bar plot in the upper panel shows the number of solutions found for each combination of three fixed points, while the bars of the right count the number of individual fixed points, regardless of their combinations. B) Phase space plots of the four most predominant combinations of fixed points. The black points indicate the random values used for initializing the simulation.

Next, we explored whether the reproduction of empirical observables depended on the different combinations of fixed points in the local dynamics. Figure 5 presents all pairs of local dynamics that significantly differed in the goodness of fit according to multiple metrics (synchronization, metastability, and Kolmogorov-Smirnov distance between distributions of FCD values). For each pair, we computed a distribution of effect size estimates (difference between the medians of both groups, Fig. 5, bottom panels) following a bootstrap procedure (see methods section), and selected as significant those pairs of dynamics for which the confidence interval (CI) of the effect size distribution did not include zero (i.e., equal medians) with a 95% confidence level. Furthermore, to filter the cases with the most significant effect size we just kept those where the lower (upper) bound of the CI was at least at distance of 0.05 from the zero. From these results, it is clear that local dynamics including stable spirals systematically outperformed unstable spirals.

**Figure 5.**
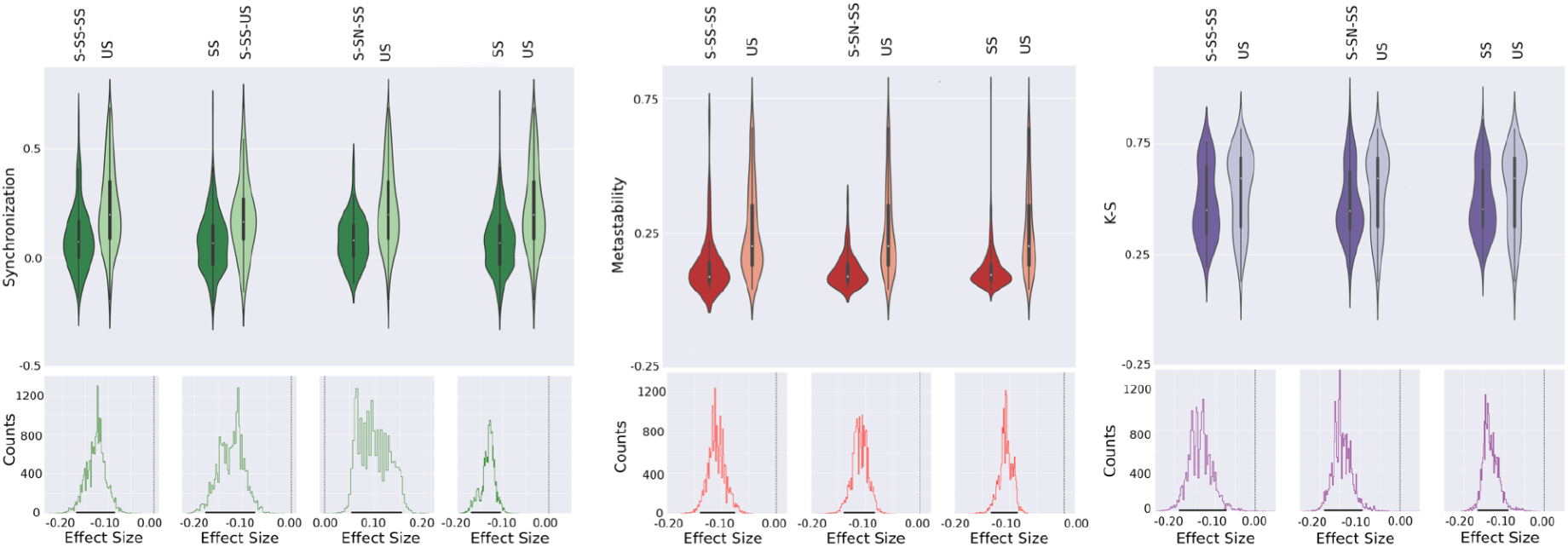
Local dynamics with stable spirals resulted in better reproduction of the empirical data in terms of synchronization, metastability, and Kolmogorov-Smirnov distance between distributions of FCD values. Violin plots present the distribution of performance metrics for all solutions with the local dynamics indicated by the labels. The bottom panels show the distribution of effect sizes obtained using bootstrap. The vertical line indicates zero, i.e., null effect size, while the 95% confidence intervals are indicated using thick black lines in the x-axis.

In the case of local dynamics with more than one stable spiral, the solution was asymptotically attracted to one of the spirals. Taking the combination S-SS-SS as an example, we found that the stable spiral with the best goodness of fit in terms of 1-SSIM was the one with the lowest absolute value of the real eigenvalue, i.e. the stable spiral with eigenvalue closest to zero outperformed all other fixed points. This was a general result valid for all combinations of fixed points and all goodness of fit metrics (Fig. 6), indicating that the best local dynamics were close to a change in stability, from unstable to stable spirals and vice versa.

**Figure 6.**
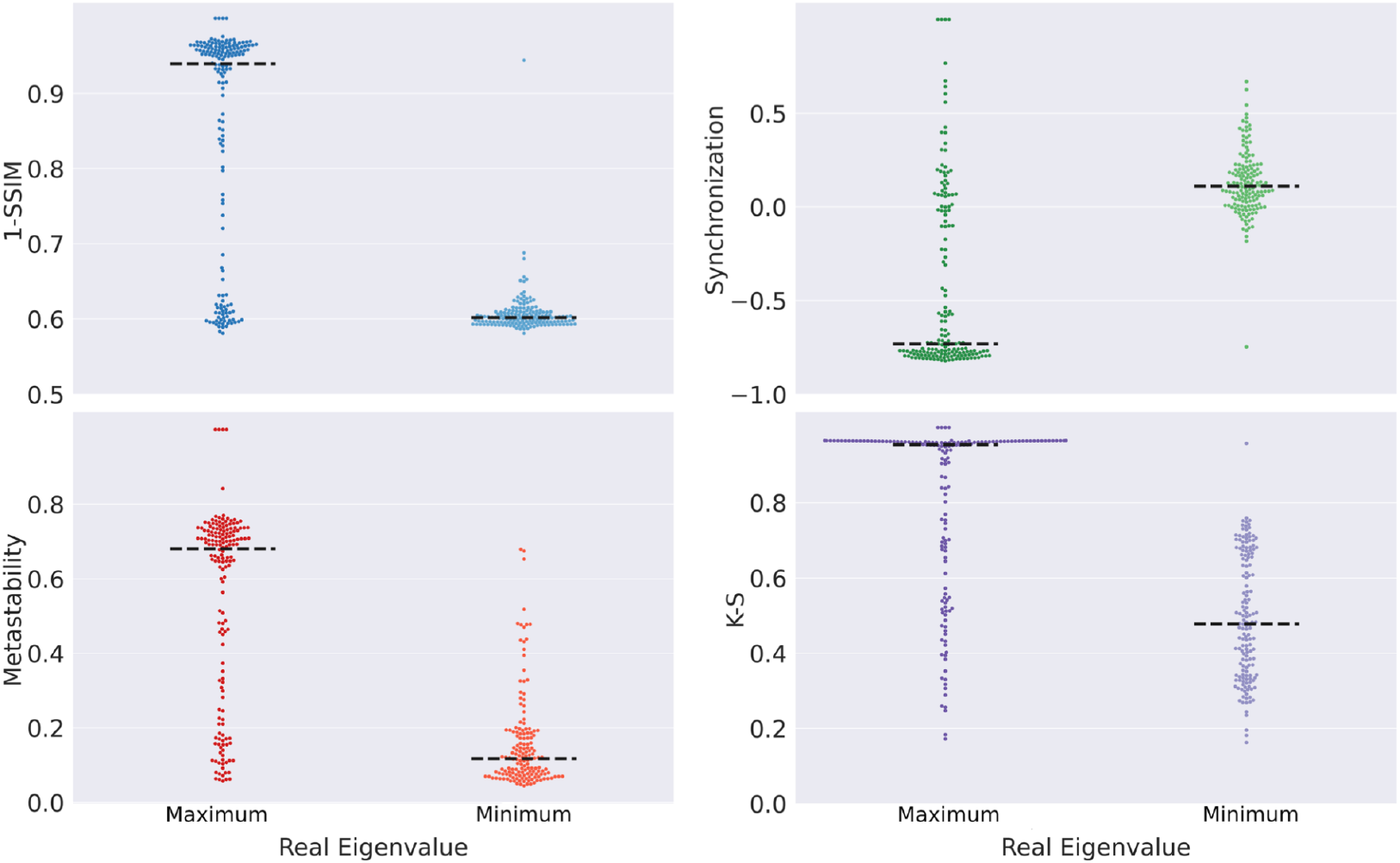
Spiral fixed points with real eigenvalues closer to zero resulted in a better reproduction of the empirical observables regardless of the type of fixed point. Comparison of the goodness of fit in terms of four different metrics for the spirals with maximum vs. minimum real eigenvalue. The black dashed lines denote the median of each distribution.

Whenever unstable spirals were present, local dynamics always were attracted to a stable limit cycle, corresponding to a periodic oscillatory behavior. It is important to note that these oscillations were not necessarily harmonic, due to the presence of non-linearities in the equations. We investigated whether departures from harmonic oscillations improved the fit to the experimental data using the same metrics as in the previous analyses. To obtain a measure of harmonicity, we obtained the time series for the optimal solutions which included a stable limit cycle; next, we converted these time series to the Fourier space and computed the spectral content relative to the dominant frequency; i.e. the whole spectrum was divided by the power of the dominant frequency. Afterwards we summed the power of all the spectrum. Thus, a highly harmonic time series concentrates most of the spectral power in the dominant frequency, resulting in total power near one; conversely, high values of the sum corresponds to anharmonic time series where the spectral power is spread across multiple frequencies. We considered oscillatory solutions corresponding to the top and bottom quartile of the harmonicity distribution and computed all goodness of fit metrics, with results presented in Fig. 7. Examples of harmonic and anharmonic oscillatory local dynamics are shown in Fig. 7A. The violin plots in Fig. 7B summarize the distribution of the performance metrics for all solutions presenting stable limit cycles of low and high anharmonicity. Using a bootstrap procedure (Fig. 7C) we showed that harmonic and anharmonic oscillatory were comparable in terms of 1-SSIM; however, harmonic solutions improved the goodness of fit with respect to synchronization and metastability, while anharmonic solutions improved the reproduction of the FCD.

**Figure 7.**
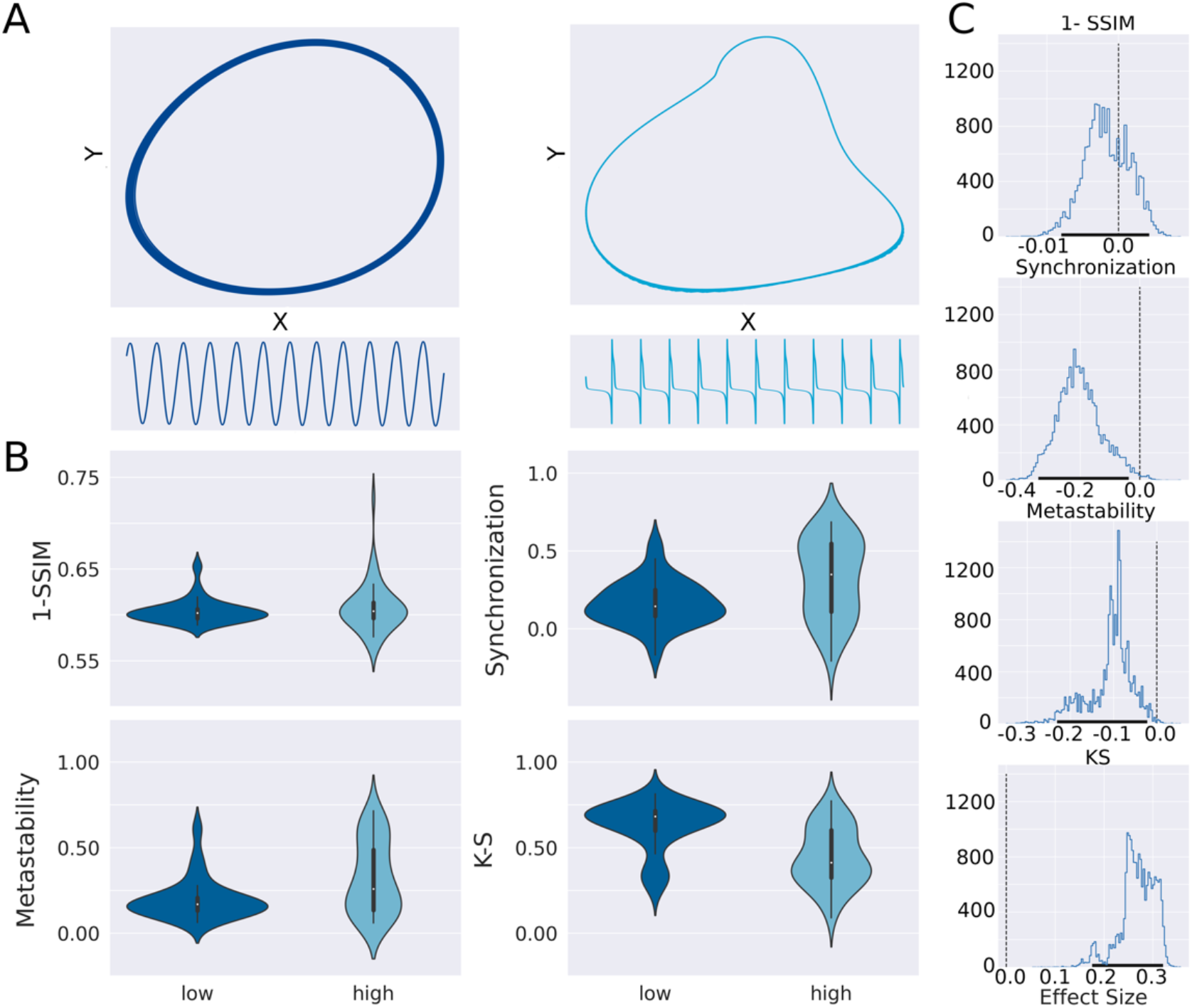
The anharmonicity of stable limit cycles in the local dynamics influenced the goodness of fit according to different metrics. A) Examples of stable limit cycles and time series of high (right) and low (left) anharmonicity. B) Violin plots summarizing the distribution of the performance metrics for all solutions presenting stable limit cycles of low and high anharmonicity. C. Distribution of effect sizes for the difference in the performance metrics obtained using bootstrap. The vertical line indicates zero, i.e., null effect size, while the 95% confidence intervals are indicated using thick black lines in the x-axis.

As a final analysis, we tested whether the optimal local dynamics inferred using our method depended on the global brain state of the participants. For this purpose, we used fMRI data acquired in the same scanner and conditions as the wakefulness data, but with participants undergoing deep sleep (n3 sleep). In previous work, a simple phenomenological model (Stuart-Landau oscillators, corresponding to the normal mode of a Hopf bifurcation) was fitted to data acquired during deep sleep, showing increased stability (i.e., distance from the bifurcation) compared to wakefulness^24^. Thus, we hypothesized that the optimal canonical dynamics inferred from deep sleep would consist of stable spirals with real eigenvalues larger than those found for wakefulness.

The results of this analysis are shown in Fig. 8. Panel A shows that, as for wakefulness, local dynamics predominantly presented one fixed point. Moreover, the most likely fixed point consisted of stable spirals, with a larger preference for these dynamics relative to wakefulness (Fig. 8B), tested for significance using a chi-squared test (p<0.001). Also, the distance to the empirical data (1-SSIM) was larger for n3 sleep compared to wakefulness, indicating more difficulty to properly capture the functional connectivity matrix. This result is consistent with previous publications applying whole-brain computational models to the same dataset^26^. Finally, a bootstrap procedure was used to compare the stable spiral real eigenvalues obtained for both conditions (Fig. 8D). We found that the real eigenvalues were within 95% confidence level in the range [-0.017, -0.013] and [-0.0149, -0.0089] for n3 sleep and wakefulness respectively, indicating a significant shift towards more negative real eigenvalues for n3, consistent with previous research^24^.

**Figure 8.**
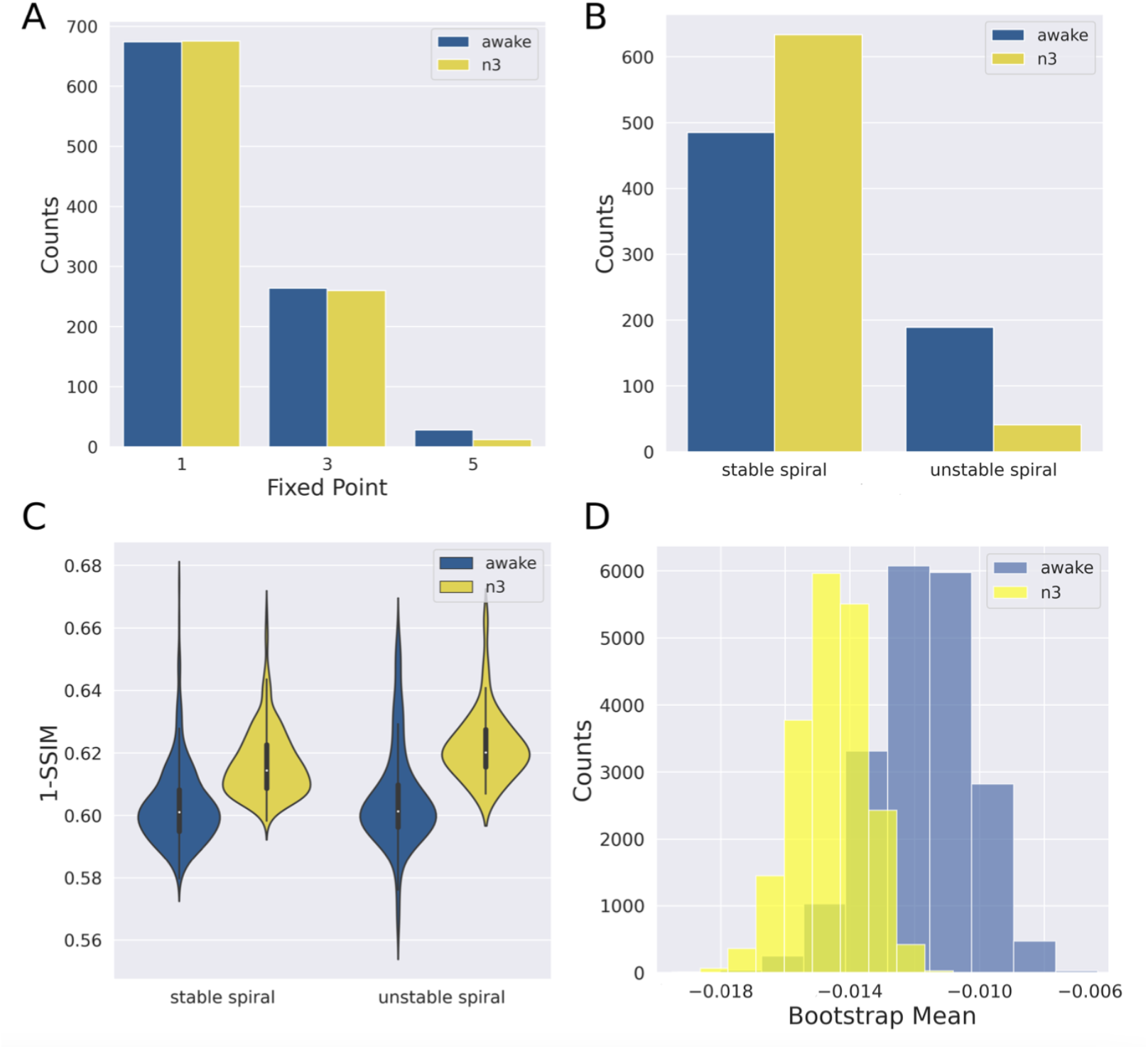
Deep sleep resulted in the stabilization of fMRI dynamics. A) Number of fixed points in the optimal local dynamics, for wakefulness and n3. B) Relative prevalence of stable spirals vs. unstable spirals for both brain states. C) Distance to the empirical data (1-SSIM) for wakefulness and n3 sleep. D) Histogram of the real eigenvalues of the stable spiral fixed points computed using a bootstrap procedure and for both brain states. The shift towards left for n3 indicates increased stability of the local dynamics relative to wakefulness.

## Discussion

We addressed a central problem in computational neuroscience: what kind of local dynamics suffice to capture the emergent macroscopic activity of the brain? A long-standing line of research has attacked this problem from a bottom-up perspective, assembling detailed biophysical descriptions of individual neurons and then characterizing the dynamical repertoire of the resulting neural mass equations^7^. While fruitful, this approach depends on the particular details of how each model is constructed and implemented, and thus leaves the possibility open that different dynamics may improve the characterization of empirical observables. We adopted the novel yet complementary top-bottom approach of exhaustively exploring a large space of possible equations for the local dynamics, focusing afterwards on characterizing the most frequently represented behaviors. We identified these local dynamics as *canonical*, in the sense that they should be included in computational models of large-scale activity to adequately reproduce empirical observables, regardless of the level of biophysical realism. Thus, our approach is useful to delineate the dynamics that should be included in whole-brain models, and thus to constraint the development of more sophisticated biophysical models.

A corollary of our results is that neurobiological realism can only improve the fit to empirical neuroimaging observables insofar certain types of local dynamics are included in the model. Also, our analysis revealed that neither the number of fixed-points nor the precise combination of fixed-points appearing in the phase space are important factors to determine the performance of a whole-brain model. Instead, local dynamics should unfold in the proximity of a specific type of attractor and also near a qualitative change in the space near that attractor (known as a bifurcation). Even though the inclusion of additive noise introduced variability in the local dynamics found by the optimization algorithm, we found that stable spirals were overrepresented in the optimal solutions. Moreover, as shown in Fig. 3, the imaginary eigenvalues of the stable spirals could take a wide range of positive values, while the real eigenvalues were predominantly negative and close to zero. This result not only indicates that the local dynamics preferentially consist of damped oscillations (stable spiral attractor), but also that the system is posed close to a bifurcation (change in the sign of the real eigenvalue). This observation is supported by the findings shown in Fig. 6, when there are multiple spirals in the solutions the best values of multiple goodness of fit metrics are obtained for spirals with real eigenvalues closer to zero. When noise-driven dynamics are close to a Hopf bifurcation, a phenomenon known as noise-induced multistability can result in the intermittent displacement between dynamical regimes (i.e. across the bifurcation)^23,32^. Thus, even if dynamics unfold in the proximity of a stable spiral attractor, the amplitude of the oscillations might not decrease steadily over time; instead, the presence of additive noise is capable of changing the nature of the solutions, giving rise to complex modulations in the amplitude of the oscillations^33^.

Some behaviors are a priori ruled out by considerations of biological plausibility; for instance, dynamics should unfold within a bounded region of phase space. Yet within these constraints, many possible scenarios were also ruled out by our analysis. Even though noise-driven linear dynamics (multivariate Ornstein-Uhlenbeck processes) are included within the space of possible models we explored^34^, our results point towards the importance of nonlinearities in the local dynamics of whole-brain models. Bistable dynamics (or other solutions given by connected saddle nodes) were also ruled out by our analysis^35-37^. Oscillations are ubiquitous in the emergent macroscopic activity of the brain, yet only those in the damped regime predominated among the optimal equations for the local dynamics. This result agrees with empirical results as well as with the dynamical repertoire of multiple models of large-scale brain activity, which feature transitions towards stable spirals through different bifurcations^22,38-40^. Finally, in the case of oscillatory dynamics (stable limit cycle), the presence of anharmonicities influenced the goodness of fit metrics, with departures from sinusoidal waveforms benefiting the reproduction of empirical FCD.

The improved performance of local dynamics with small real eigenvalues highlights the importance of the proximity to a bifurcation. Also, this suggests that the presence of a Hopf bifurcation (i.e. transition between noisy and oscillatory dynamics) is required to capture multiple independent observables derived from fMRI data, regardless of the biophysical sophistication of the model. Accordingly, phenomenological whole-brain models including this type of bifurcation have been used in recent years to simulate different physiological and pathological brain states, as well as to study in silico their behavior under multiple forms of external perturbations^13,24-27^. Thus, our results can be interpreted as a hypothesis-free validation of the Hopf model (also known as Stuart-Landau oscillator), although in our case the oscillations were not always harmonic. Future research should explore whether certain deviations from harmonic oscillations are required to improve the description of macroscopic brain activity, as has already been supported by experiments^41^.

It is important to note that we only explored local dynamics described by two variables, one interpreted as a direct readout of the recorded signal and the other necessary as an auxiliary variable to increase the diversity of behaviors displayed by the model. Including a third variable would open the possibility of deterministic chaos in the equations, which could represent an alternative to noise-induced metastability to generate complex modulations of the oscillatory dynamics. Recently, we showed that deterministic chaos can favor the simultaneous reproduction of multiple neuroimaging observables at the same time, as it “stretches” the range where complex oscillations are produced, in contrast to the fine tuning of parameters necessary for noise-induced metastability^42^. Moreover, chaos and noise could be complementary, as their combination might enhance the dynamical repertoire of whole-brain models, endowing them with desirable properties for the reproduction of empirical data^43^. Future research should incorporate a third variable to the analysis, thus allowing to investigate the relative importance of chaos and noise-driven metastability in a data-driven way.

Our results also corroborated that the optimal local dynamics depend on the global brain state. We investigated differences in the parameters found for wakefulness and n3 (deep) sleep. While the optimal number of fixed-points did not change between conditions, we found that stable spirals became more predominant during sleep. Consistently, we also found a shift towards negative real eigenvalues, indicative of the stabilization of the local dynamics during unconsciousness, as suggested by multiple experimental reports^44,45^. In particular, this is consistent with a previous study showing the same result for a model based on Stuart-Landau oscillators^24^; however, our result should be considered more general as it was found by analyzing a much larger set of possible dynamics, without a priori constraining the solutions to be near a Hopf bifurcation.

In summary, we developed a top-bottom characterization of the canonical dynamics that should be included in whole-brain activity models to adequately capture empirical observables. Future work should address the implications of these dynamics in terms of large-scale information processing associated with behavior and cognitive function, extending our results towards other model organisms and imaging modalities, and incorporating our findings to the process of constructing and validating biophysically realistic models of macroscopic brain activity.

## Materials and methods

### Data availability

fMRI time series used to construct the empirical observables can be found in the following link: https://doi.org/10.6084/m9.figshare.20372172.v1

### Participants and EEG-fMRI data acquisitions

A cohort of 63 healthy subjects participated in the data acquisition protocol (36 females, mean ± SD age of 23.4 ± 3.3 years). Written informed consent was obtained from all subjects. The experimental protocol was approved by the ethics committee of Goethe-Universität Frankfurt, Germany (protocol number: 305/07). The subjects were reimbursed for their participation. All experiments were conducted in accordance with the relevant guidelines and regulations, and the Declaration of Helsinki. Participants were scanned for 50 minutes using previously published acquisition parameters. For the analysis of awake subjects, we selected a subgroup of 9 participants who did not fall asleep throughout the complete duration of the scan (confirmed by assessment of the simultaneous EEG according to standard sleep staging rules). In this way, we obtained long fMRI recordings with the purpose of robustly estimating observables related to functional connectivity dynamics.

### fMRI data processing

Using Statistical Parametric Mapping (SPM8, www.fil.ion.ucl.ac.uk/spm), raw fMRI data were realigned, normalized and spatially smoothed using a Gaussian kernel with 8 mm^3^ full width at half maximum. Data was then re-sampled to 4 × 4 × 4 mm resolution. Note that re-sampling introduced local averaging of blood-oxygen-level-dependent (BOLD) signals, which were eventually averaged over larger cortical and sub-cortical regions of interest as determined by the automatic anatomic labeling (AAL) atlas^46^. Data was denoised by regressing out cardiac, respiratory and residual motion time series estimated with the RETROICOR method, and then band-pass filtered in the 0.01–0.1 Hz range using a sixth order Butterworth filter^47,48^.

### Anatomical connectivity matrix

The anatomical connectivity matrix was obtained applying diffusion tensor imaging (DTI) to diffusion weighted imaging (DWI) recordings from 16 healthy right-handed participants (11 men and 5 women, mean age: 24.75 ± 2.54 years) recruited online at Aarhus University, Denmark. Subjects with psychiatric or neurological disorders (or a history thereof) were excluded from participation. We refer to a previous publication for details of the MRI acquisition parameters.

Anatomical connectivity networks were constructed following a three-step process. First, the regions of the whole-brain network were defined using the AAL atlas. Second, the connections between nodes in the whole-brain network (edges) were estimated applying probabilistic tractography to the DTI data obtained for each participant. Third, results were averaged across participants. DTI preprocessing was performed using the probtrackx tool of the FSL diffusion imaging toolbox (Fdt) (www.fsl.fmrib.ox.ac.uk/fsl/fslwiki/FDT) with default parameters. Next, the local probability distributions of fiber directions were estimated at each voxel. The connectivity probability from a seed voxel i to another voxel j was defined as the proportion of fibers passing through voxel i that reached voxel j, sampling a total of 5000 streamlines per voxel. This was extended from the voxel to the region level, with each region of interest consisting of n voxels, so that 5000 × n fibers were sampled. The connectivity probability from region i to region j was calculated as the number of sampled fibers in region i that connected the two regions, divided by 5000 × n, where n represents the number of voxels in region i. The resulting anatomical connectivity matrices were thresholded at 0.1% (i.e. a minimum of five streamlines), resulting in the anatomical connectivity matrices used as coupling in the whole-brain models.

### Whole-brain model construction

Following previous work^27^, we constructed computational models of whole-brain activity by assigning local dynamical rules to 90 nodes spanning the whole cortical and subcortical grey matter. These nodes were coupled using an anatomical connectivity matrix C_n,s_ which contained in its n, s entry an estimate of the number of white matter tracts connecting nodes n and s (see previous section). We introduced a parameter G to globally scale the C_n,s_ matrix, thus modeling changes in the overall strength of inter-areal coupling.

The fMRI signal corresponding to node n was simulated by the variable x_n_, obtained from the differential equation modeling the local dynamics of that node, integrated using a Euler-Maruyama algorithm with a time step of 0.1. For each parameter combination, we computed observables (see below) by averaging a total of 30 independent simulations. Simulated time series were down-sampled to match the sampling frequency of the fMRI data.

### Local dynamics ansatz

We consider a general ansatz for the noise-driven local dynamics of node n, given by polynomial equations on variables x and y up to degree five,

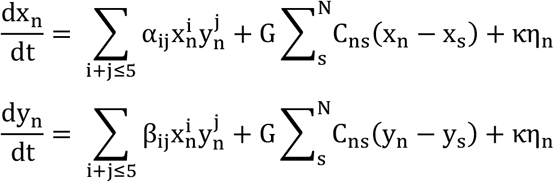

Here, η_n_(t) corresponds to additive Gaussian noise at node n scaled by parameter κ, C_ns_ is the anatomical coupling matrix scaled by parameter G, and α_ij_, β_ij_ are the coefficients of the polynomial terms which determine the nature of the local dynamics. The choice of polynomial terms follows from the objective of determining the optimal canonical local dynamics, since it is known that systems close to a bifurcation are topologically equivalent to a normal form, which can be written as a polynomial^29^.

### Genetic algorithm for parameter optimization

The genetic algorithm started with a generation of 10 sets of parameters (“individuals”) chosen uniformly at random in the range [-0.15, 0.15] for each of the 42 parameters. A score proportional to the target function was assigned to each individual. Afterwards, a group of individuals was chosen based on their score (“parents”). Operations of crossover between parents generate new possible solutions (“offspring”). Mutation and elite selection were applied to create the next generation of solutions. These three operations can be briefly described as follows: 1) elite selection occurs when an individual of a generation shows an extraordinarily low target function (i.e., high goodness of fit) in comparison to the other individuals, thus this solution is replicated without changes in the next generation; 2) the crossover operator consists of combining two selected parents to obtain a new individual that carries information from each parent to the next generation; 3) the mutation operator can change an individual of the offspring set to induce a random alteration in any of its parameters.

Following previous work^27^, 20% of each new generation was created by elite selection and 80% by crossover of the parents, with a 5% chance of possible mutations in the offspring group. Each generation was used iteratively as the seed for the next generation until 125 generations were created. This criterion was chosen to guarantee convergence of solutions. After applying the optimization algorithm, the parameter values corresponding to the best fit were used to explore the phase space of local dynamics (see subsection “Fixed-point analysis and classification”).

### Target fMRI observables and goodness of fit metrics

We obtained the functional connectivity matrix by computing the Pearson correlation coefficient between fMRI signals (empirical or simulated) at all pairs of regions in the parcellation. This resulted in a symmetric matrix whose i, j entry contained the correlation between the signal extracted from regions i and j. To measure the similarity between empirical and simulated matrix we used the structural similarity index (SSIM)^30^, a metric which combines the similarity in terms of the Euclidean and correlation distances (for further details see previous implementations^27^)

To characterize the time-dependent structure of resting state fluctuations we computed the functional connectivity dynamics (FCD) matrix^25^. Using 148 sliding 60 s sliding windows with 40 s overlap, we calculated the temporal evolution of the functional connectivity, and then obtained the t_1_, t_2_ entry of the 148 × 148 symmetric FCD matrix by computing the Pearson correlation coefficient between the upper triangular part of functional connectivity matrices at times t_1_ and t_2_. The similarity between empirical and simulated FCD matrices was given by the Kolmogorov-Smirnov distance (maximum difference between the cumulative distribution functions of the two samples) between the upper diagonal part of the corresponding matrices.

To compute the synchronization and metastability^31^, we first extracted the phases of the bandpass filtered fMRI signals from each of the 90 regions and for each subject, and then obtained the analytic narrowband signal, a(t) = x(t) + iH[x(t)], where i is the imaginary unit, x(t) the original signal, and H[x(t)] its Hilbert transform. The instantaneous phase was then obtained as ϕ(t) = arg(a(t)). We computed the Kuramoto order parameter, R(t), as:

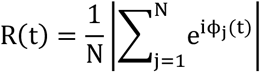

In this equation, N is the total number of nodes and ϕ_j_(t) represents the instantaneous phase of node j. The order parameter R(t) measures the instantaneous global phase synchrony of the system, ranging from 0 (absence of synchrony) to 1 (full synchronization). The temporal average and standard deviation of R(t) represent the synchronization and metastability, respectively. The first of these two metrics indicates the global and temporally averaged degree of synchronization between all the nodes in the system, whereas the second gives information about temporal variability in the level of synchronization. For both observables, we compared empirical and simulated data by subtracting them and normalizing by the value obtained for the empirical data.

### Parameter optimization

We used a stochastic optimization method (genetic algorithm) to determine the optimal 42 parameters (α_ij_, β_ij_, i + j ≤ 5) of the local dynamics to maximize the SSIM between empirical and simulated functional connectivity matrices. The global coupling scaling parameter was fixed at G=0.5, as determined previously^27^. After optimization, we computed multiple observables to compare the empirical and simulated time series.

### Fixed-point analysis and classification

The parameter optimization procedure was repeated 1000 times, and for each set of optimal parameters we investigated the asymptotic behavior of the resulting equations for the local dynamics. First, we determined the fixed-points, i.e., the points of invariance of the dynamics, by introducing a grid in the range x, y ∈ [−100,100] and searching the roots of the equations for 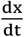 and 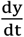. Once the fixed points were found, we classified them according to the following criteria.

Let J be the Jacobian matrix of the optimal model parameters:

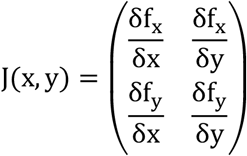

where f_x_, f_y_ are the equations for 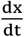 and 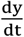, respectively. We obtained one 2×2 matrix by computing J(x, y) at each fixed-point. The stability at each of these points follow from the eigenvalues of the Jacobian. In the two-dimensional case, stability can also be computed from the trace (τ) and determinant (Δ), and the results can be classified as follows^49^:

1. If Δ ≤0, the fixed point is a saddle (attracts the dynamics along one direction, while repelling it along another)
2. If Δ >0, τ <0 and 4Δ – τ^2^ >0, the fixed point is a stable spiral (damped oscillations)
3. If Δ >0, τ <0 and 4Δ – τ^2^ <0, the fixed point is a stable node (attracts the dynamics from all directions)
4. If Δ >0, τ >0 and 4Δ – τ^2^ >0, the fixed point is an unstable spiral (oscillations with increasing amplitude)
5. If Δ >0, τ >0 and 4Δ – τ^2^ <0, the fixed point is an unstable node (repels the dynamics from all directions)

After all fixed-points and their stability were computed, we re-simulated the dynamics with initial conditions close to each of these points and computed the target fMRI observables and their associated goodness of fit (see “Target fMRI observables and goodness of fit metrics” subsection). This procedure was repeated 30 times for each fixed-point and the resulting goodness of fit metrics were averaged across all iterations.

### Effect size and bootstrapping

We obtained effect size estimates between two groups of values by computing the difference between the medians of both groups, since the values do not necessarily follow a normal distribution. We used bootstrapping to obtain a distribution of effect sizes estimates, allowing to determine confidence interval (CI) of the effect size distributions and thus whether an overlap exists at 95% confidence level.

## Acknowledgments

This work was supported by grants PICT-2019-02294 (Agencia I+D+i, Argentina) and ANID/FONDECYT Regular 1220995 (Chile).

## Notes

### Competing Interest Statement

The authors have declared no competing interest.

## References

1 Chialvo, D. R. Emergent complex neural dynamics. Nature Physics 6, 744–750 (2010).

2 Bassett, D. S. & Gazzaniga, M. S. Understanding complexity in the human brain. Trends Cogn Sci 15, 200–209, doi:10.1016/j.tics.2011.03.006 (2011).

3 Tononi, G. & Edelman, G. M. Consciousness and complexity. Science 282, 1846–1851, doi:10.1126/science.282.5395.1846 (1998).

4 Bullmore, E. & Sporns, O. Complex brain networks: graph theoretical analysis of structural and functional systems. Nat Rev Neurosci 10, 186–198, doi:10.1038/nrn2575 (2009).

5 Sporns, O. Network attributes for segregation and integration in the human brain. Curr Opin Neurobiol 23, 162–171, doi:10.1016/j.conb.2012.11.015 (2013).

6 Cofre, R. et al. Whole-Brain Models to Explore Altered States of Consciousness from the Bottom Up. Brain Sci 10, doi:10.3390/brainsci10090626 (2020).

7 Deco, G., Jirsa, V. K., Robinson, P. A., Breakspear, M. & Friston, K. The dynamic brain: from spiking neurons to neural masses and cortical fields. PLoS Comput Biol 4, e1000092, doi:10.1371/journal.pcbi.1000092 (2008).

8 Murray, J. D., Demirtas, M. & Anticevic, A. Biophysical Modeling of Large-Scale Brain Dynamics and Applications for Computational Psychiatry. Biol Psychiatry Cogn Neurosci Neuroimaging 3, 777–787, doi:10.1016/j.bpsc.2018.07.004 (2018).

9 Mejias, J. F. & Wang, X. J. Mechanisms of distributed working memory in a large-scale network of macaque neocortex. Elife 11, doi:10.7554/eLife.72136 (2022).

10 Jirsa, V. K. et al. The Virtual Epileptic Patient: Individualized whole-brain models of epilepsy spread. Neuroimage 145, 377–388, doi:10.1016/j.neuroimage.2016.04.049 (2017).

11 Deco, G. & Kringelbach, M. L. Great expectations: using whole-brain computational connectomics for understanding neuropsychiatric disorders. Neuron 84, 892–905, doi:10.1016/j.neuron.2014.08.034 (2014).

12 Yonatan Sanz Perl, C. P., Ignacio Pérez Ipiña, Athena Demertzi, Vincent Bonhomme, Charlotte Martial, Rajanikant Panda, Jitka Annen, Agustin Ibañez, Morten Kringelbach, Gustavo Deco, Helmut Laufs, Jacobo Sitt, Steven Laureys, Enzo Tagliazucchi Perturbations in dynamical models of whole-brain activity dissociate between the level and stability of consciousness. Plos Computational Biology (2021).

13 Perl, Y. S. et al. Generative Embeddings of Brain Collective Dynamics Using Variational Autoencoders. Phys Rev Lett 125, 238101, doi:10.1103/PhysRevLett.125.238101 (2020).

14 Arbabyazd, L. et al. Virtual Connectomic Datasets in Alzheimer’s Disease and Aging Using Whole-Brain Network Dynamics Modelling. eNeuro 8, doi:10.1523/ENEURO.0475-20.2021 (2021).

15 Perl, Y. S., Pallavicini, C., Ipiña, I. P., Kringelbach, M., Deco, G., Laufs, H., Tagliazucchi, E. Data augmentation based on dynamical systems for the classification of brain states.. Chaos, Solitons and Fractals 139 (2020).

16 Falcon, M. I., Jirsa, V. & Solodkin, A. A new neuroinformatics approach to personalized medicine in neurology: The Virtual Brain. Curr Opin Neurol 29, 429–436, doi:10.1097/WCO.0000000000000344 (2016).

17 Deco, G., Jirsa, V., McIntosh, A. R., Sporns, O. & Kotter, R. Key role of coupling, delay, and noise in resting brain fluctuations. Proc Natl Acad Sci U S A 106, 10302–10307, doi:10.1073/pnas.0901831106 (2009).

18 Deco, G. et al. Whole-Brain Multimodal Neuroimaging Model Using Serotonin Receptor Maps Explains Non-linear Functional Effects of LSD. Curr Biol 28, 3065–3074 e3066, doi:10.1016/j.cub.2018.07.083 (2018).

19 Kringelbach, M. L. et al. Dynamic coupling of whole-brain neuronal and neurotransmitter systems. Proc Natl Acad Sci U S A 117, 9566–9576, doi:10.1073/pnas.1921475117 (2020).

20 Vohryzek, J., Cabral, J., Vuust, P., Deco, G. & Kringelbach, M. L. Understanding brain states across spacetime informed by whole-brain modelling. Philos Trans A Math Phys Eng Sci 380, 20210247, doi:10.1098/rsta.2021.0247 (2022).

21 Shine, J. M., Aburn, M. J., Breakspear, M. & Poldrack, R. A. The modulation of neural gain facilitates a transition between functional segregation and integration in the brain. Elife 7, doi:10.7554/eLife.31130 (2018).

22 Schirner, M., Kong, X., Yeo, B. T. T., Deco, G. & Ritter, P. Dynamic primitives of brain network interaction. Neuroimage 250, 118928, doi:10.1016/j.neuroimage.2022.118928 (2022).

23 Deco, G., Jirsa, V. K. & McIntosh, A. R. Emerging concepts for the dynamical organization of resting-state activity in the brain. Nat Rev Neurosci 12, 43–56, doi:10.1038/nrn2961 (2011).

24 Jobst, B. M. et al. Increased Stability and Breakdown of Brain Effective Connectivity During Slow-Wave Sleep: Mechanistic Insights from Whole-Brain Computational Modelling. Sci Rep 7, 4634, doi:10.1038/s41598-017-04522-x (2017).

25 Deco, G., Kringelbach, M. L., Jirsa, V. K. & Ritter, P. The dynamics of resting fluctuations in the brain: metastability and its dynamical cortical core. Sci Rep 7, 3095, doi:10.1038/s41598-017-03073-5 (2017).

26 Sanz Perl, Y. et al. Perturbations in dynamical models of whole-brain activity dissociate between the level and stability of consciousness. PLoS Comput Biol 17, e1009139, doi:10.1371/journal.pcbi.1009139 (2021).

27 Ipina, I. P. et al. Modeling regional changes in dynamic stability during sleep and wakefulness. Neuroimage 215, 116833, doi:10.1016/j.neuroimage.2020.116833 (2020).

28 Brunton, S. L., Proctor, J. L. & Kutz, J. N. Discovering governing equations from data by sparse identification of nonlinear dynamical systems. Proc Natl Acad Sci U S A 113, 3932–3937, doi:10.1073/pnas.1517384113 (2016).

29 Murdock, J. Normal forms. Scholarpedia 1 (2006).

30 W. Zhou, A. C. B. H.R. Sheikh, E.P. Simoncelli. Image qualifty assessment: from error visibility to structural similarity. IEEE Trans. Image Process 13, 600–612 (2004).

31 Acebrón, J., Bonilla, L. L., Pérez Vicente, C. J., Ritort, F., Spigler, R. The Kuramoto model: A simple paradigm for synchronization phenomena. Reviews of modern physics 77, 137 (2005).

32 Ghosh, A., Rho, Y., McIntosh, A. R., Kotter, R. & Jirsa, V. K. Noise during rest enables the exploration of the brain’s dynamic repertoire. PLoS Comput Biol 4, e1000196, doi:10.1371/journal.pcbi.1000196 (2008).

33 Juel, A., Darbyshire, A. G., Mullin, T. The effect of noise on pitchfork and Hopf bifurcations. Proceedings of the Royal Society of London (series A) 453, 2627–2647 (1997).

34 Saggio, M. L., Ritter, P. & Jirsa, V. K. Analytical Operations Relate Structural and Functional Connectivity in the Brain. PLoS One 11, e0157292, doi:10.1371/journal.pone.0157292 (2016).

35 Freyer, F., Roberts, J. A., Ritter, P. & Breakspear, M. A canonical model of multistability and scale-invariance in biological systems. PLoS Comput Biol 8, e1002634, doi:10.1371/journal.pcbi.1002634 (2012).

36 Freyer, F. et al. Biophysical mechanisms of multistability in resting-state cortical rhythms. J Neurosci 31, 6353–6361, doi:10.1523/JNEUROSCI.6693-10.2011 (2011).

37 Buendia, V., Di Santo, S., Villegas, P., Burioni, R., Muñoz, M. Self-organized bistability and its possible relevance for brain dynamics. Physical Review Research 013318 (2020).

38 Spyropoulos, G. et al. Spontaneous variability in gamma dynamics described by a damped harmonic oscillator driven by noise. Nat Commun 13, 2019, doi:10.1038/s41467-022-29674-x (2022).

39 Galinsky, V. L. & Frank, L. R. Universal theory of brain waves: from linear loops to nonlinear synchronized spiking and collective brain rhythms. Phys Rev Res 2, doi:10.1103/PhysRevResearch.2.023061 (2020).

40 Hutcheon, B. & Yarom, Y. Resonance, oscillation and the intrinsic frequency preferences of neurons. Trends Neurosci 23, 216–222, doi:10.1016/s0166-2236(00)01547-2 (2000).

41 Cole, S. R. & Voytek, B. Brain Oscillations and the Importance of Waveform Shape. Trends Cogn Sci 21, 137–149, doi:10.1016/j.tics.2016.12.008 (2017).

42 Piccinini, J. et al. Noise-driven multistability vs deterministic chaos in phenomenological semi-empirical models of whole-brain activity. Chaos 31, 023127, doi:10.1063/5.0025543 (2021).

43 Orio, P. et al. Chaos versus noise as drivers of multistability in neural networks. Chaos 28, 106321, doi:10.1063/1.5043447 (2018).

44 Massimini, M. et al. Breakdown of cortical effective connectivity during sleep. Science 309, 2228–2232, doi:10.1126/science.1117256 (2005).

45 Solovey, G. et al. Loss of Consciousness Is Associated with Stabilization of Cortical Activity. J Neurosci 35, 10866–10877, doi:10.1523/JNEUROSCI.4895-14.2015 (2015).

46 Tzourio-Mazoyer, N. et al. Automated anatomical labeling of activations in SPM using a macroscopic anatomical parcellation of the MNI MRI single-subject brain. Neuroimage 15, 273–289, doi:10.1006/nimg.2001.0978 (2002).

47 Glover, G. H., Li, T. Q. & Ress, D. Image-based method for retrospective correction of physiological motion effects in fMRI: RETROICOR. Magn Reson Med 44, 162–167, doi:10.1002/1522-2594(200007)44:1<162::aid-mrm23>3.0.co;2-e (2000).

48 Cordes, D. et al. Frequencies contributing to functional connectivity in the cerebral cortex in “resting-state” data. AJNR Am J Neuroradiol 22, 1326–1333 (2001).

49 Shnol, E. E. Stability of equilibria. Scholarpedia 2 (2007).

